# When the Background Matters: Topic-Dependent reference lists in GWAS and Exome Analyses

**DOI:** 10.64898/2026.08.14.744838

**Authors:** Brian Timoney, Paolo Guasoni, Komal Zade, Snow Bach, Daniela Tropea

## Abstract

Gene Ontology (GO) Biological Process overrepresentation analysis is widely used to interpret gene lists from genetic studies, yet results depend critically on the background (universe/reference list) against which enrichment is tested. This paper examines how genome–exome background mismatch alters GO Biological Process significance and induces annotation-driven bias. First, Monte Carlo simulations across multiple input gene list sizes show that enrichment p-values shift systematically when lists sampled from an exome-like universe are tested against a genome background (and vice versa), producing both inflation and deflation of significance depending on GO term composition; these shifts increase with gene list size. Second, applied analyses of gene lists derived from Genome-Wide Association Studies (GWAS) and Whole Exome Studies (WES) across brain, immune, and metabolic domains demonstrate that background choice changes the set of significant GO IDs, yielding reference-specific terms consistent with both Type I errors (false positives) and Type II errors (false negatives). Because genome backgrounds are commonly used by default, the practical risk is greatest when WES-derived lists are analyzed with genome reference lists. To support reproducible best practice, we provide a simple command set for selecting and documenting study-appropriate backgrounds and for assessing sensitivity of GO Biological Process results to the chosen universe.

## Introduction

High-throughput genetic and molecular profiling routinely produces lists of genes associated with a phenotype or contrast of interest, including hits from genome-wide association studies (GWAS), whole-exome sequencing (WES), and transcriptomic studies. Although such lists are informative, interpreting them gene-by-gene often provides limited insight into the cellular mechanisms that differ between conditions. Therefore, highly accessible platforms have been developed to organize input genes into an output where genes are grouped according to biological processes, pathways, or other parameters. The grouping is based on curated annotations, whereby – based on scientific publication-a gene is attributed a specific function or participation in a biological process. The annotations are then used in the functional enrichment analysis: a process that determines how much the input list is over-represented for curated annotations.

One of the most commonly used tool is Gene Ontology (GO)(2), which highlights biological processes (but also cellular components and molecular functions) that may underlie the observed signal. Other popular tools (DAVID, Enrichr, g:Profiler, clusterProfiler, etc.) use the same principle.

A critical but frequently under-specified component of overrepresentation analysis is the choice of the *background* gene universe (also referred to as the reference list). Conceptually, enrichment asks whether genes annotated to a given GO class, for example “Biological Process” occur in the input list more often than expected by chance under random sampling from the background. Because the expected overlap depends on the composition of the background, using an inappropriate universe can shift enrichment p-values and, consequently, the set of processes reported as significant. Thus, choosing an appropriate background is essential.

The GO Consortium explicitly advises using as background “the list of all genes from which your smaller analysis list was selected”. In practice, however, this step is frequently overlooked. *Wijesooriya et al. (2022)* surveyed 186 papers and found that 95% of GO overrepresentation analysis (ORA) analyses either used an inappropriate background or failed to report it. This widespread neglect is not just a procedural detail—it can profoundly bias results. When the background is mis-specified, the null model of enrichment is wrong, leading to systematic inflation or deflation of p-values and misleading sets of “enriched” GO terms.

The majority of tools have the default option to use all genes in the genome as the background. This option is often defensible for GWAS-derived gene lists, which originate from genome-wide scans, but it is generally misaligned with assays that interrogate a restricted subset of genes. For WES, the effective universe is largely protein-coding genes. For next-generation sequencing transcriptomic studies, the appropriate background depends on how RNA molecules are selected and captured during library preparation and what technique has been used for transcript quantification. For microarray studies, the background should reflect the probes present on the array used for measurement. Treating these restricted assays as if they were sampled from the full genome risks introducing systematic bias in downstream functional interpretation (Reimand et al., 2019; Mubeen et al., 2022). An RNA-seq experiment effectively tests only the genes expressed above detection limits. In such cases, using all ∼60,000 human genes (including non-coding or unassayed genes) as the background introduces *unrelated genes with no chance of appearing in the list, thereby producing an artificial enrichment*.

There are several other possible biases in enrichment studies. Here, we focus on GO Biological Process annotations and quantify the consequences of genome–exome background mismatch due to an inadequate reference list. We use simulation-based experiments that compare enrichment outcomes from random gene lists sampled from a genome universe versus an exome universe, evaluated under both genome and exome backgrounds. To isolate the effect of background choice as directly as possible, we emphasize GWAS- and WES-style gene lists rather than expression-based inputs, thereby avoiding additional pitfalls that are common in transcriptomic data.

We then provide applied demonstrations using gene lists drawn from studies spanning three biological domains—brain, immune, and metabolism—to assess whether background-driven distortions are consistent across contexts. Across simulations and real gene lists, we show that an incorrect background can alter GO Biological Process significance in both directions, creating potential Type I errors (false positives) and Type II errors (false negatives), with effects that vary across domains.

Finally, we provide ready-to-use R code to define study-appropriate backgrounds and to compare results against the default genome background, supporting transparent reporting and sensitivity analyses in routine GO Biological Process enrichment workflows (code to be made available on GitHub upon acceptance).

## Methods

### Monte Carlo Simulations

#### Reference List Creation

Throughout, we use the term **background** (also referred to as the *universe* or *reference list*) to denote the set of genes against which GO Biological Process overrepresentation is tested.

We constructed both a genome background and an exome background from the org.Hs.egGO2ALLEGS database, which provides mappings between GO identifiers and Entrez Gene identifiers annotated to each term or its child terms in the GO hierarchy. For Biological Process analyses, we retrieved all unique genes associated with the “BP” ontology to define the **genome background** (i.e., the background typically used by enrichGO in the clusterProfiler package).

To derive the **exome background** for simulation, we mapped Entrez Gene identifiers to *protein* Ensembl identifiers using the MyGene.info package. Genes with missing/NULL protein Ensembl mappings were removed; the remaining genes defined the exome background.

#### Random Sampling (Input Gene Lists)

Random input gene lists were generated by sampling from either the genome background or the exome background. Using the base R sample() function, we drew 40,000 independent random gene lists for each input size in {10, 20, 50, 100, 150, 200, 250}. We included smaller sizes (10 and 20) because many available exome derived study gene lists were short, and we included larger sizes up to 250 to reflect common practice in molecular profiling pipelines; for example, GEO2R on the NCBI website uses a default maximum of 250 significant genes for transcriptomic studies. We used 40,000 iterations per condition to obtain stable empirical estimates of significance rates across GO terms.

#### Biological Process Enrichment Analysis

To quantify the effect of background choice on GO Biological Process enrichment, we performed overrepresentation analysis (hypergeometric test) on random input gene lists under different background configurations. Specifically, we evaluated the following combinations:

1. **Genome input samples with Genome Reference**
2. **Genome input samples with Exome Reference**
3. **Exome input samples with Genome Reference**
4. **Exome input samples with Exome Reference**

For efficiency and reproducibility, we implemented a local version of the enrichGO procedure that stores the background sets and their associated GO gene sets locally (i.e., without server calls) and returns p-values only. For each input list, the function outputs one p-value per GO ID, representing the significance of overlap between the input genes and the GO term gene set under the hypergeometric test.

Consistent with enrichGO defaults, we restricted analyses to GO IDs with gene set sizes between **10 and 500** to limit extreme sparsity and reduce dependence among tested terms. However, unlike the default enrichGO output, we did **not** remove GO IDs with p-value = 1, because our simulation analyses require a complete p-value vector for every GO ID in every iteration.

For each GO ID and each input/background configuration, we calculated the **proportion of simulations** (out of 40,000) in which that GO ID was significant at **α = 0.05**. We then generated two summary comparisons: (i) Fig. 1 compares exome-sampled versus genome-sampled inputs when p-values are computed using the genome background, and (ii) Fig. 2 compares p-values computed using the genome versus exome background for inputs sampled from the exome background.

**Fig. 1.**
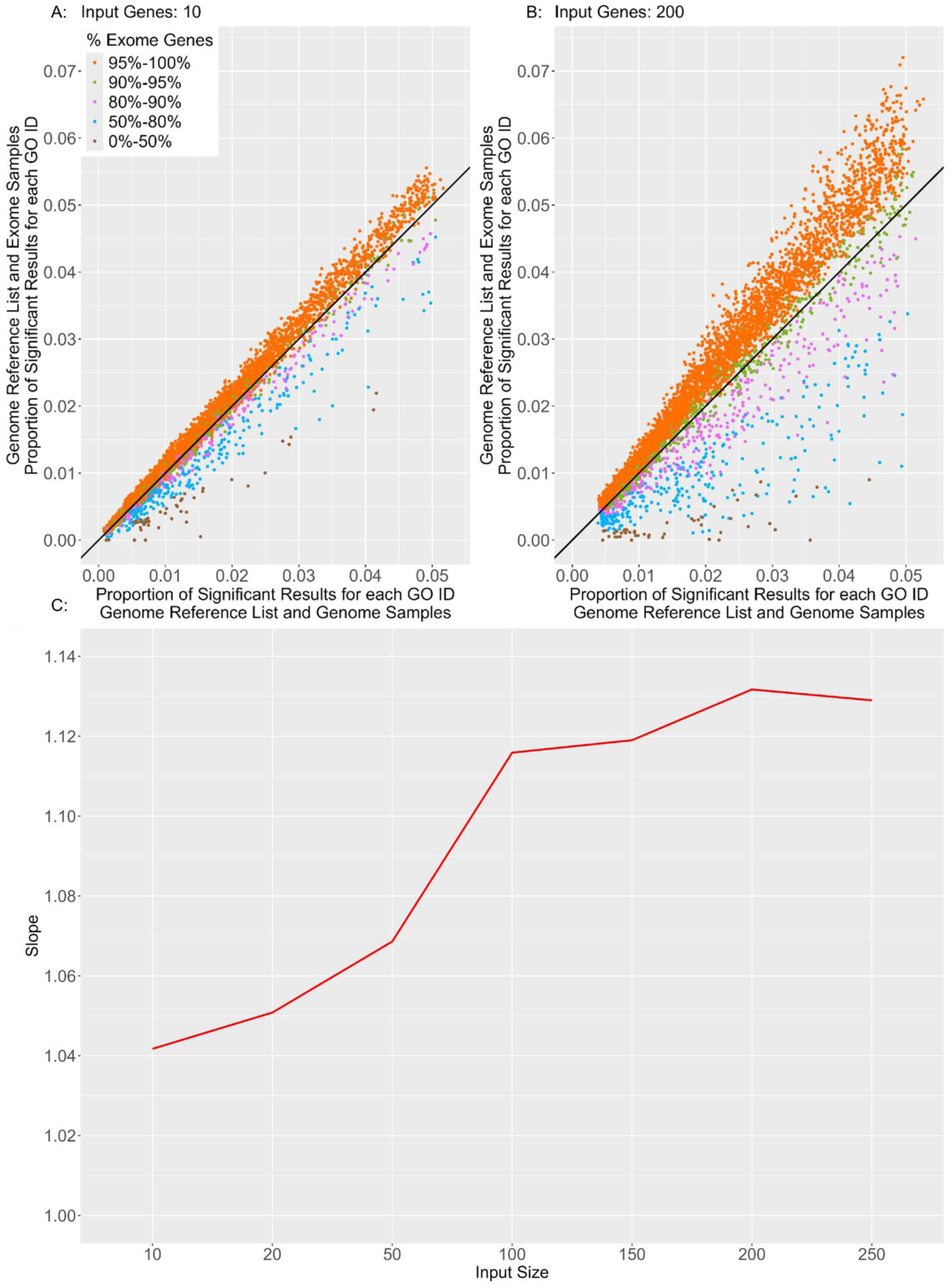
Effect of the Input Type and the Input Size on the Proportion of times each GO ID is considered significant at a 5% significance level when using the Genome Reference List. Fig. 1A and 1B show how using exome sample inputs (vertical) creates more significant results compared to using genome sample inputs (horizontal), when using the genome reference list as default. The comparison between Fig. 1A (10 genes in the input) and 1B (200 genes in the input) shows how the extra significance due to the input type increases further when using larger input sizes. If the exome sample inputs produced the same amount of significance as the genome sample inputs we would expect all of the points to lie on the black line with slope 1 (Significance produced using exome inputs = Slope x Significance produced using genome inputs). However, Fig.1C shows that as we increase our input size (horizontal), the significance produced using exome sample inputs heavily outweighs the significance produced when using genome sample inputs (i.e. the slope increases significantly - vertical). The increase in significance when using exome sample inputs highlights that only a small proportion of the genes related to GO IDs between sizes 10 and 500, are genes which do not code for proteins. Put differently, the majority of the genes annotated in GO IDs are coding for proteins, causing an annotation bias in the GO analysis when the reference list is not adequate (Gaudet & Dessimoz, 2017).

**Fig. 2.**
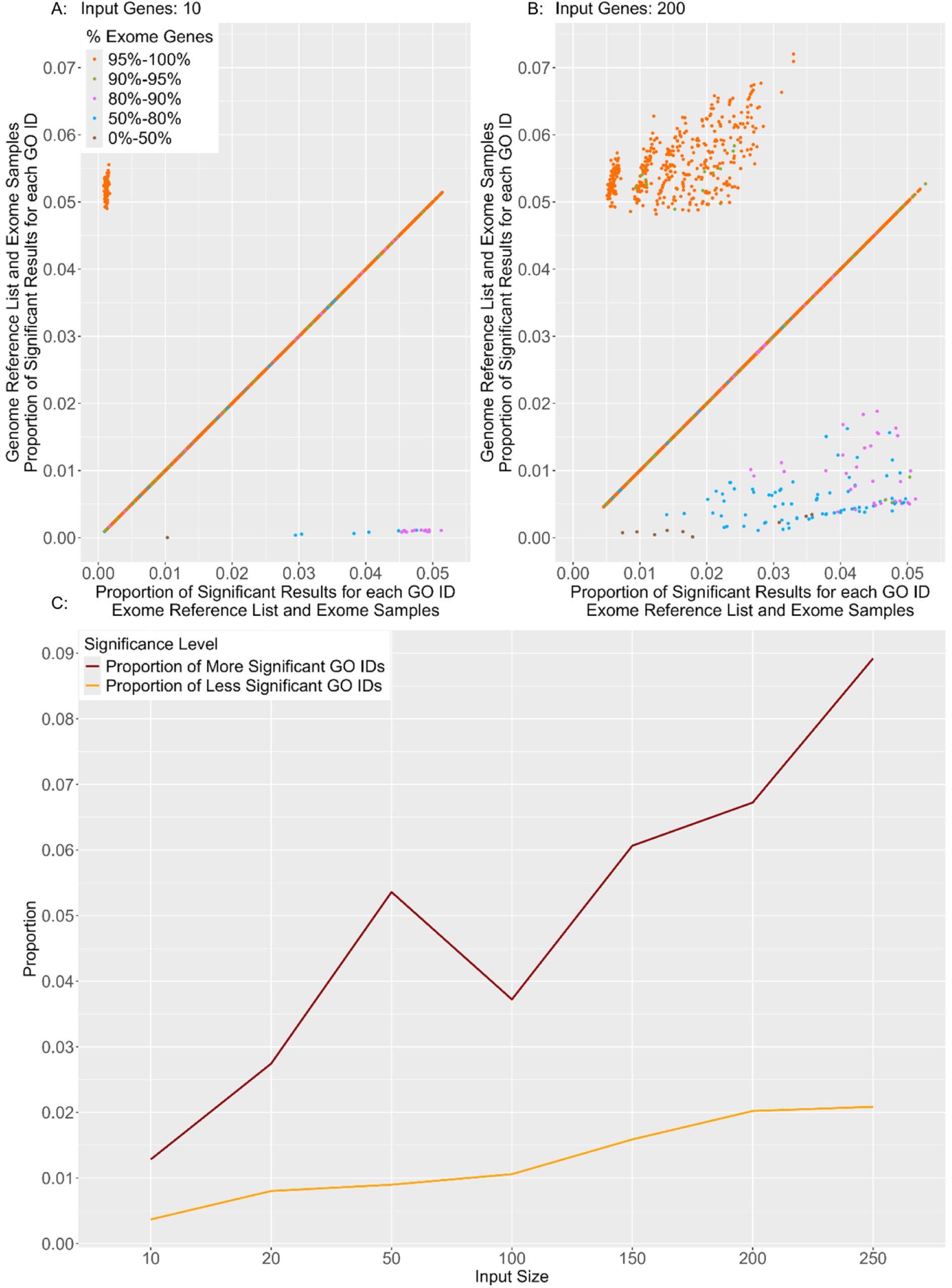
Effect of background choice and input size on GO Biological Process significance rates for exome-sampled inputs. For each GO identifier (GO ID), the proportion of Monte Carlo iterations in which the term was significant at α = 0.05 is shown for exome-sampled input lists under two backgrounds: genome background (vertical) and exome background (horizontal). (A–B) Scatterplots for input sizes of 10 genes (A) and 200 genes (B); the black diagonal indicates equal significance rates under the two backgrounds. Points above the diagonal represent GO IDs that are significant more often under the genome background (inflation relative to the exome background), whereas points below represent GO IDs that are significant less often under the genome background (deflation). (C) The proportion of GO IDs above and below the diagonal varies with input size, reflecting that the set of GO IDs exhibiting inflation or deflation depends on input size, GO ID size, and the fraction of exome genes represented in each GO ID; the trend is not strictly monotone across sizes. Because the GO ID sets associated with the genome and exome backgrounds are not identical, comparisons in (A–B) are restricted to GO IDs present under both backgrounds.

Finally, because GO IDs available under the genome background are not identical to those available under the exome background (due to background-specific term membership and term-size filtering), comparisons between genome- and exome-background results (Fig. 2) were restricted to GO IDs **present in both sets**.

### Selection and Processing of GWAS and WES Datasets

To assess how background selection influences real study outputs, we analyzed published gene lists from Genome-Wide Association Studies (GWAS) and whole exome sequencing (WES) studies spanning three biological domains: **Brain**, **Immune**, and **Metabolism**. Relevant studies were identified using keyword searches for “GWAS”, “exome”, “brain”, “immune”, and “metabolism”. Each study was assigned a unique code, and Table 1 catalogues study details, including PubMed IDs (PMIDs).

**Table 1.** List of GWAS and WES studies for analysing background selection effects. Table catalogues the published GWAS and WES studies across three biological domains (Brain, Immune, Metabolism), selected to verify the impact of background selection on real study outputs. Each domain includes 6 GWAS and 3 WES studies. Columns indicate: biological domain, study type (GWAS/WES), unique study ID, and corresponding PubMed ID for the study.

| Domain | Study type | Study ID | PMIDs |
| --- | --- | --- | --- |
| Brain | GWAS | BG1 | 24162737 |
| Brain | GWAS | BG2 | 24126926 |
| Brain | GWAS | BG3 | 35379992 |
| Brain | GWAS | BG4 | 35396580 |
| Brain | GWAS | BG5 | 21614001 |
| Brain | GWAS | BG6 | 24842889 |
| Brain | WES | BE1 | 29181857 |
| Brain | WES | BE2 | 25363760 |
| Brain | WES | BE3 | 24410847 |
| Immune | GWAS | IG1 | 32296059 |
| Immune | GWAS | IG2 | 28425483 |
| Immune | GWAS | IG3 | 33676448 |
| Immune | GWAS | IG4 | 38368726 |
| Immune | GWAS | IG5 | 36750564 |
| Immune | GWAS | IG6 | 24076602 |
| Immune | WES | IE1 | 27111861 |
| Immune | WES | IE2 | 24001973 |
| Immune | WES | IE3 | 34539730 |
| Metabolism | GWAS | MG1 | 26068415 |
| Metabolism | GWAS | MG2 | 26308950 |
| Metabolism | GWAS | MG3 | 31287004 |
| Metabolism | GWAS | MG4 | 38201907 |
| Metabolism | GWAS | MG5 | 29973135 |
| Metabolism | GWAS | MG6 | 23545492 |
| Metabolism | WES | ME1 | 37560457 |
| Metabolism | WES | ME2 | 35568032 |
| Metabolism | WES | ME3 | 32778825 |

Gene lists were obtained primarily from the main text or supplementary materials. When a study did not provide a gene list directly, genes were extracted from relevant GO databases based on study details indexed in PubMed, and these study-associated gene lists were used for downstream analyses.

All gene identifiers were converted to **Entrez IDs** prior to enrichment. Using customized R code, we performed GO Biological Process enrichment under both the genome and exome backgrounds, enabling direct comparison of background-dependent outputs for a given input list. Enrichment outputs were organized into (i) GO IDs obtained using the genome background, (ii) GO IDs obtained using the exome background, (iii) GO IDs common to both backgrounds, and background-specific terms (unique genome GO IDs; unique exome GO IDs). These categories were used to highlight GO IDs detected under one background but not the other.

We then examined overlaps among enriched GO IDs (within and across Brain, Immune, and Metabolism domains), comparing GWAS and WES outputs within domains and assessing cross-domain overlap for GWAS-only, WES-only, and combined datasets. Recurrent GO IDs across multiple studies were noted alongside their known biological functions and pathways. Potential false positives—defined here as **unique genome GO IDs detected from WES gene lists** or **unique exome GO IDs detected from GWAS gene lists**—were used to illustrate how background choice can alter the biological interpretation of study-derived gene lists.

## Results

### Pitfalls of an Incorrect Reference List

We first examined how the choice of **background** (also referred to as the universe/reference list) affects the enrichment of GO Biological Process using simulations under controlled sampling and GO IDs with sizes between 10 and 500. The GO annotation set is dominated by protein-coding (exome) genes. Consistent with this annotation bias, exome derived input lists generate systematically more significant GO IDs than genome-derived input lists when both are evaluated against the **genome background** (Fig.1). The identity line identifies the GO IDs which have the same chance of being significant when GWAS or WES studies are analyzed using the genome as a reference list. The divergence from the identity line increases with larger input gene list sizes, indicating that the inflationary effect of exome-like inputs becomes more pronounced as list size grows.

We then fixed the input to exome derived gene lists and compared enrichment results obtained using the **genome background** versus the **exome background** (Fig.2). This comparison shows two opposing effects: some GO IDs become significant more often under the genome background (inflation relative to the exome background), whereas others become significant less often (deflation). In practical terms, an incorrect background can therefore introduce both **Type I errors (false positives)** and **Type II errors (false negatives)**, depending on the composition of each GO ID. The extent of divergence increases with input size, but the relationship is not strictly monotone across all sizes because the set of GO IDs that can shift toward “more significant” depends jointly on input size, GO ID size, and the proportion of exome genes represented in the background. Because the GO ID sets available under genome and exome backgrounds are not identical, genome-versus-exome background comparisons are restricted to GO IDs present in both backgrounds (Fig.2).

Overall, the simulation results demonstrate that the default option of using a genome background can systematically distort GO Biological Process enrichment for exome derived inputs, and that the direction of distortion is GO-ID-specific.

### Verifying the Impact of Reference list Selection on published GWAS and Exome Datasets

To evaluate whether the simulation patterns are reflected in real study outputs, we compiled gene lists from six GWAS and three WES studies spanning three biological domains: brain (Bellenguez et al., 2022; Kenny et al., 2014; Lambert et al., 2013; Trubetskoy et al., 2022; Vacic et al., 2014; Voineagu et al., 2011, Cukier et al., 2014; De Rubeis et al., 2014; Patel et al., 2018), immune (Beecham et al., 2013; Han et al., 2020; Khunsriraksakul et al., 2023; Li et al., 2021; Liu et al., 2024; Qiu et al., 2017, Dinwiddie et al., 2013; Liu et al., 2021; Mak et al., 2016), and metabolism (Draisma et al., 2015; Ho et al., 2023; Jiao et al., 2015; Lalonde et al., 2019; Naik et al., 2013; Rimpelä et al., 2018, Adhikari et al., 2020; Bomba et al., 2022; Ferreira et al., 2023) (Table 1). From each study gene list, twenty genes were randomly extracted and analyzed using our R workflow under both genome and exome backgrounds. For each dataset, enriched GO IDs were partitioned into: (i) **common GO IDs** (significant under both backgrounds), (ii) **unique genome GO IDs** (significant only under the genome background), and (iii) **unique exome GO IDs** (significant only under the exome background).

When considering the significant GO IDs across domains, we found that GWAS datasets—and particularly datasets in the brain domain—yield higher counts of significant GO IDs (Fig.3). Unique genome GO IDs were observed across all domains, whereas unique exome GO IDs were comparatively uncommon, and exome studies generally showed fewer common GO IDs shared across backgrounds. Despite matching the number of studies per domain, the fraction of unique exome GO IDs detected in exome datasets remained low (Fig.3), indicating that background choice can change the composition of reported GO Biological Process results even when the input lists are held to the same size.

**Fig. 3.**
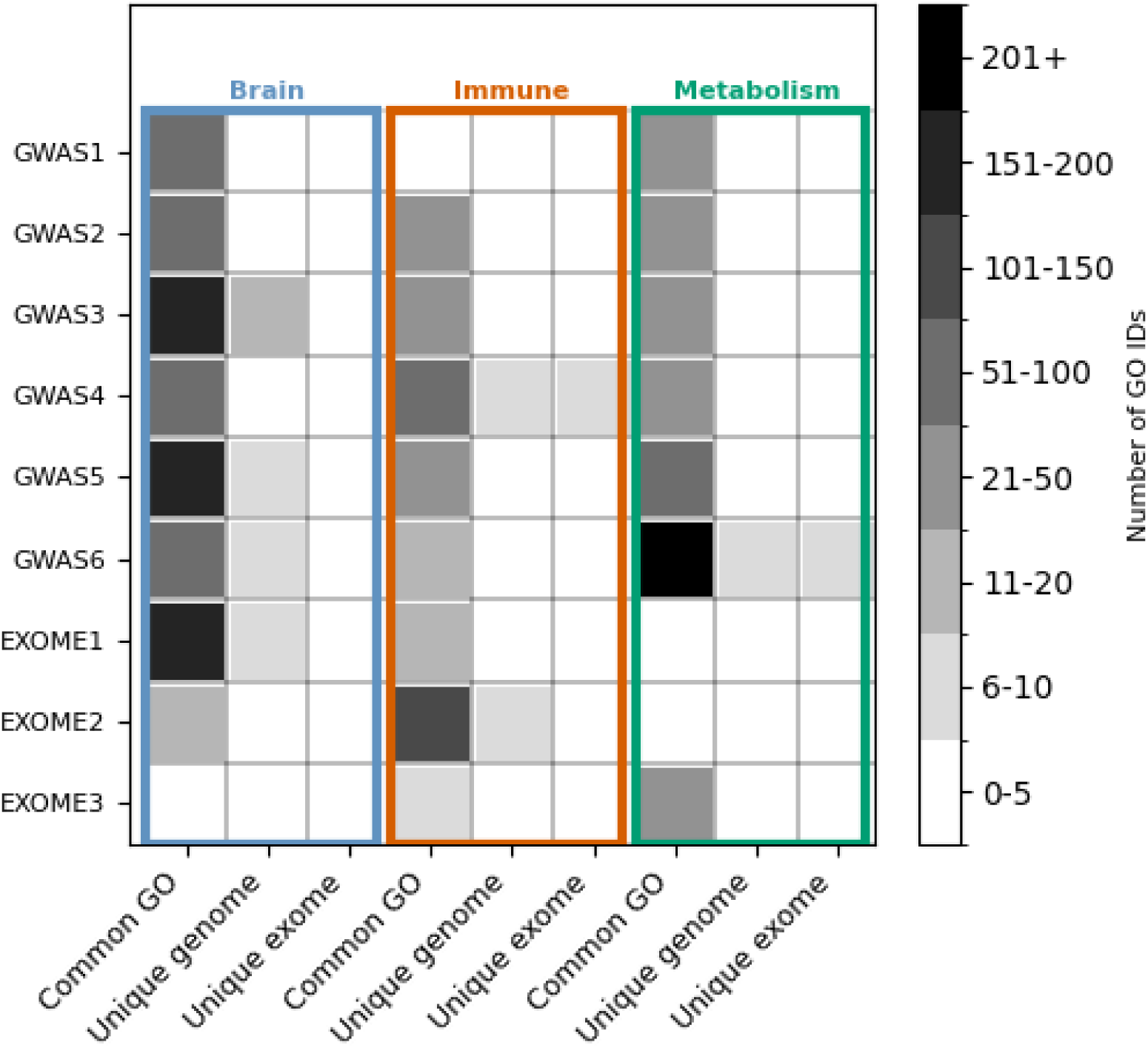
Distribution of Gene Ontology (GO) ID counts by study type and biological domains. The horizontal axis groups studies by biological domain (brain, immune, metabolism), and the vertical axis arranges studies by study type, with whole-exome sequencing (WES) studies at the bottom and genome-wide association studies (GWAS) at the top. Within each study, three bars summarize the number of significant GO IDs: common to genome and exome backgrounds (left), unique to the genome background (middle), and unique to the exome background (right). Shading intensity encodes GO ID counts, with lighter grey indicating lower counts and darker grey indicating higher counts.

The overlap between GWAS and WES enrichment outputs is evident in the brain and immune datasets but not in metabolism (Fig.3). This pattern suggests that consistency of enriched Biological Process signals across study types may vary by domain in the available examples, although the limited number of publicly accessible WES gene lists limits the possibility of generalizing our results.

Next, we tested whether any enriched GO IDs were shared across domains. No GO IDs were common to all three domains in either the exome-only datasets or the combined GWAS+WES analyses. This lack of cross-domain overlap is consistent with the expectation that enriched Biological Process signals—and their sensitivity to background specification—may be domain-specific in the studied sets.

Altogether, these results point to a different sensibility to the reference list for study domain.

#### Detection of False Positives from Genome and Exome Reference List

We then investigated what are the consequences of using an inadequate reference list in different study domains. To categorize the background-driven errors, we defined **potential false positives (Type I errors)** as background-specific GO IDs that appear under an implausible or mismatched background choice as follows:

1. **Unique genome GO IDs** detected when WES datasets are analyzed using the **genome background** (the most common default-use scenario).
2. **Unique exome GO IDs** detected when GWAS datasets are analyzed using the **exome background** (less likely in routine practice, but included here to illustrate symmetry of the effect).

Under these definitions, potential false positives occur across all biological domains but show no cross-domain overlap (Fig.4), suggesting that background-driven spurious enrichment depends on domain context rather than arising from a small set of universally “problematic” GO IDs. Only a small number of unique genome GO IDs are shared across domains (Table 2), consistent with the same conclusion. Immune datasets show the highest incidence of unique genome GO IDs under genome-background analysis of WES gene lists, while metabolism datasets show the lowest (Fig.4). Qualitatively, most false-positive GO IDs in immune and metabolism studies align with their expected domain pathways; in contrast, a subset of false-positive GO IDs in brain WES analyses correspond to immune and metabolic processes (Table 2), highlighting either shared biology or background-sensitive annotation structure.

**Fig. 4.**
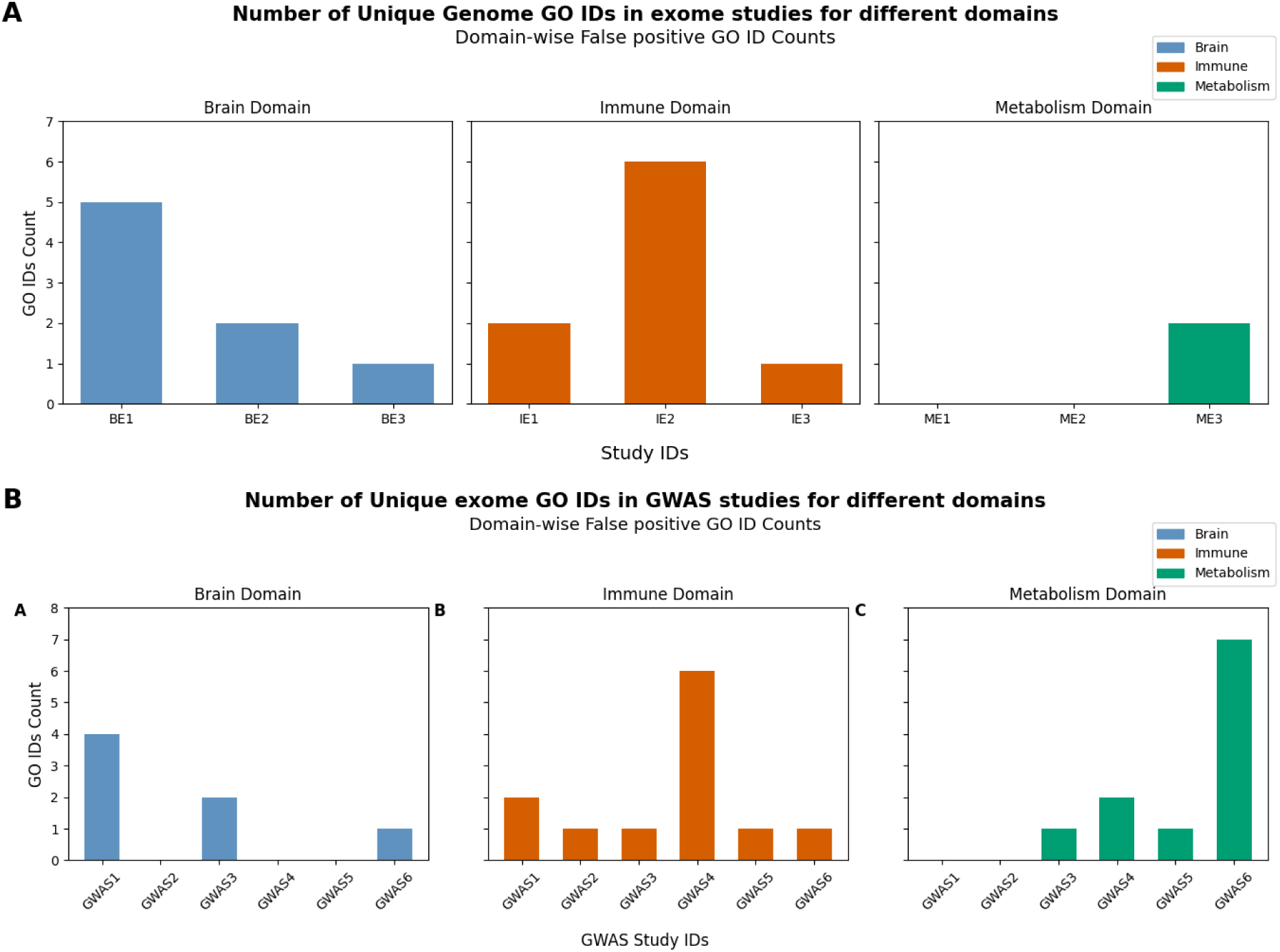
Domain-stratified counts of background-specific GO IDs used as operational false positives. (A) Counts of GO IDs significant only under the genome background (unique genome GO IDs) for whole-exome sequencing (WES) studies analyzed using the genome background. (B) Counts of GO IDs significant only under the exome background (unique exome GO IDs) for genome-wide association studies (GWAS) analyzed using the exome background. The horizontal axis groups studies by biological domain (brain, immune, metabolism) and labels individual studies by study ID; the vertical axis shows the count of background-specific GO IDs per study. Domain color coding is: brain (blue), immune (red), and metabolism (green).

**Table 2.**
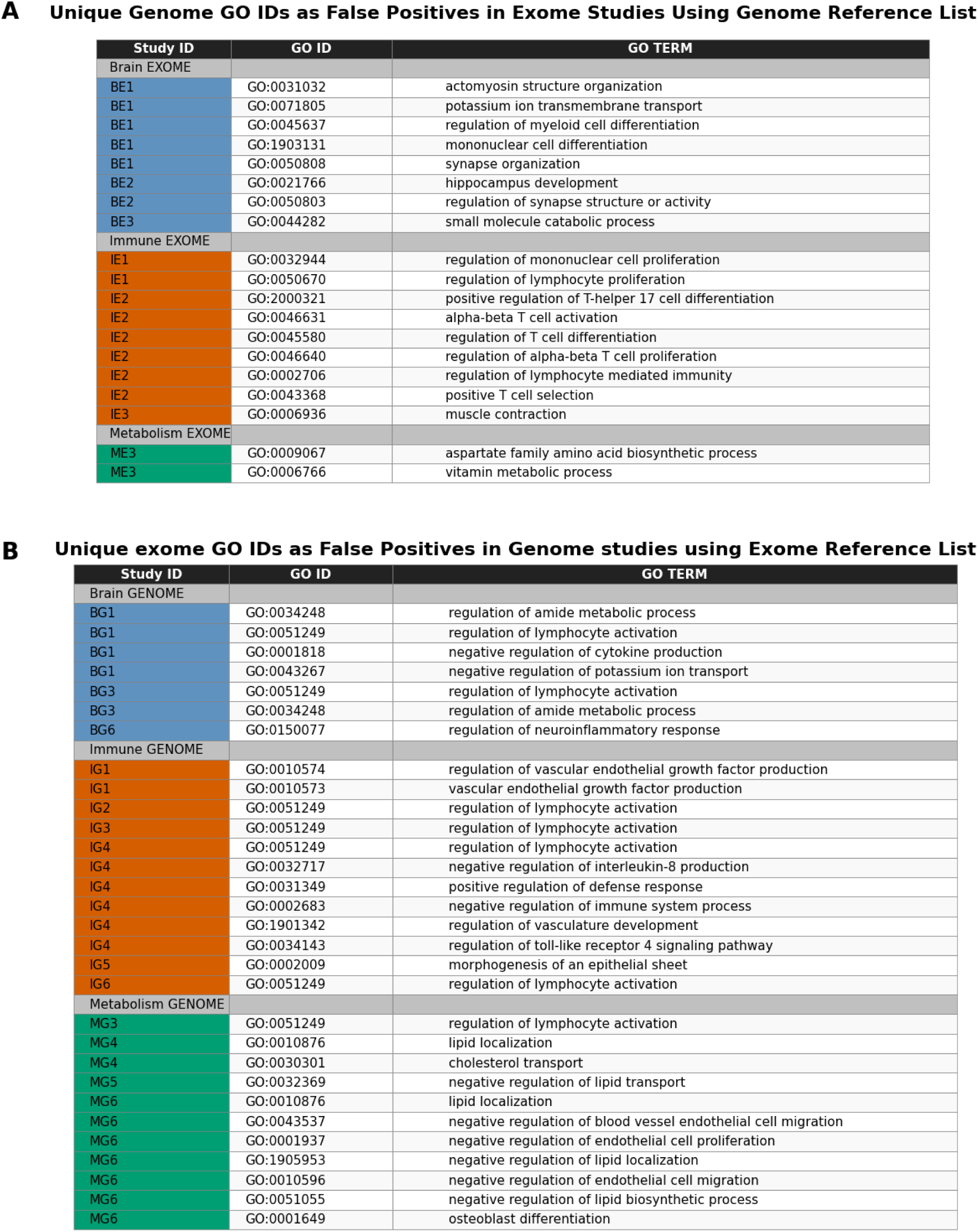
Background-specific GO Biological Process terms used as operational false positives under mismatched backgrounds. (A) GO identifiers (GO IDs) significant only under the genome background (unique genome GO IDs) for whole-exome sequencing (WES) datasets analyzed using the genome background. (B) GO IDs significant only under the exome background (unique exome GO IDs) for genome-wide association studies (GWAS) datasets analyzed using the exome background. For each entry, the table reports the study ID (with domain classification: brain, immune, metabolism), the GO ID, and the associated GO Biological Process term (as summarized in *Fig. 4*).

A Unique Genome GO IDs as False Positives in Exome Studies Using Genome Reference List
| Study ID | GO ID | GO TERM |
| --- | --- | --- |
| Brain EXOME |  |  |
| BE1 | GO:0031032 | actomyosin structure organization |
| BE1 | GO:0071805 | potassium ion transmembrane transport |
| BE1 | GO:0045637 | regulation of myeloid cell differentiation |
| BE1 | GO:1903131 | mononuclear cell differentiation |
| BE1 | GO:0050808 | synapse organization |
| BE2 | GO:0021766 | hippocampus development |
| BE2 | GO:0050803 | regulation of synapse structure or activity |
| BE3 | GO:0044282 | small molecule catabolic process |
| Immune EXOME |  |  |
| IE1 | GO:0032944 | regulation of mononuclear cell proliferation |
| IE1 | GO:0050670 | regulation of lymphocyte proliferation |
| IE2 | GO:2000321 | positive regulation of T-helper 17 cell differentiation |
| IE2 | GO:0046631 | alpha-beta T cell activation |
| IE2 | GO:0045580 | regulation of T cell differentiation |
| IE2 | GO:0046640 | regulation of alpha-beta T cell proliferation |
| IE2 | GO:0002706 | regulation of lymphocyte mediated immunity |
| IE2 | GO:0043368 | positive T cell selection |
| IE3 | GO:0006936 | muscle contraction |
| Metabolism EXOME |  |  |
| ME3 | GO:0009067 | aspartate family amino acid biosynthetic process |
| ME3 | GO:0006766 | vitamin metabolic process |

**Table 2. Background-specific GO Biological Process terms used as operational false positives under mismatched backgrounds.**
| Study ID | GO ID | GO TERM |
| --- | --- | --- |
| Brain GENOME |  |  |
| BG1 | GO:0034248 | regulation of amide metabolic process |
| BG1 | GO:0051249 | regulation of lymphocyte activation |
| BG1 | GO:0001818 | negative regulation of cytokine production |
| BG1 | GO:0043267 | negative regulation of potassium ion transport |
| BG3 | GO:0051249 | regulation of lymphocyte activation |
| BG3 | GO:0034248 | regulation of amide metabolic process |
| BG6 | GO:0150077 | regulation of neuroinflammatory response |
| Immune GENOME |  |  |
| IG1 | GO:0010574 | regulation of vascular endothelial growth factor production |
| IG1 | GO:0010573 | vascular endothelial growth factor production |
| IG2 | GO:0051249 | regulation of lymphocyte activation |
| IG3 | GO:0051249 | regulation of lymphocyte activation |
| IG4 | GO:0051249 | regulation of lymphocyte activation |
| IG4 | GO:0032717 | negative regulation of interleukin-8 production |
| IG4 | GO:0031349 | positive regulation of defense response |
| IG4 | GO:0002683 | negative regulation of immune system process |
| IG4 | GO:1901342 | regulation of vasculature development |
| IG4 | GO:0034143 | regulation of toll-like receptor 4 signaling pathway |
| IG5 | GO:0002009 | morphogenesis of an epithelial sheet |
| IG6 | GO:0051249 | regulation of lymphocyte activation |
| Metabolism GENOME |  |  |
| MG3 | GO:0051249 | regulation of lymphocyte activation |
| MG4 | GO:0010876 | lipid localization |
| MG4 | GO:0030301 | cholesterol transport |
| MG5 | GO:0032369 | negative regulation of lipid transport |
| MG6 | GO:0010876 | lipid localization |
| MG6 | GO:0043537 | negative regulation of blood vessel endothelial cell migration |
| MG6 | GO:0001937 | negative regulation of endothelial cell proliferation |
| MG6 | GO:1905953 | negative regulation of lipid localization |
| MG6 | GO:0010596 | negative regulation of endothelial cell migration |
| MG6 | GO:0051055 | negative regulation of lipid biosynthetic process |
| MG6 | GO:0001649 | osteoblast differentiation |

Conversely, **unique exome GO IDs** detected when analyzing WES datasets under the exome background represent signals that would be missed under the genome background, and thus constitute an operational set of **potential false negatives (Type II errors)** under incorrect default background choice (Table 3). Together, Tables 2 and 3 illustrate that incorrect background selection can both introduce additional significant processes and suppress processes that are significant under an assay-appropriate background.

**Table 3.** GO Biological Process terms significant only under the exome background for WES datasets. The table lists GO identifiers (GO IDs) significant only under the **exome background** (unique exome GO IDs) for whole-exome sequencing (WES) studies analyzed using the exome background. For each entry, the study ID (with domain classification: brain, immune), the GO ID, and the associated GO Biological Process term are reported.

| Study ID | GO ID | GO TERM |
| --- | --- | --- |
| **Brain EXOME** |  |  |
| BE1 | GO:0034248 | regulation of amide metabolic process |
| BE1 | GO:0002683 | negative regulation of immune system process |
| BE3 | GO:0010876 | lipid localization |
| **Immune EXOME** |  |  |
| IE1 | GO:0051249 | regulation of lymphocyte activation |
| IE2 | GO:0051249 | regulation of lymphocyte activation |
| IE2 | GO:0032660 | regulation of interleukin-17 production |
| IE2 | GO:0032620 | interleukin-17 production |

Finally, we summarized the most recurrent GO IDs by domain and study type in both GWAS and WES datasets. (Table 4). Several of the frequently identified immune GO IDs recur across both GWAS and WES datasets, consistent with the within-domain overlap observed in Fig.3, and emphasizing the robustness of immune-related enrichment driven by shared gene signatures. A direct side-by-side comparison of outputs under correct versus incorrect background selection (Fig.5) illustrates how background choice changes not only the number of significant GO IDs, but also which GO IDs are prioritized as most significant.

**Fig. 5.**
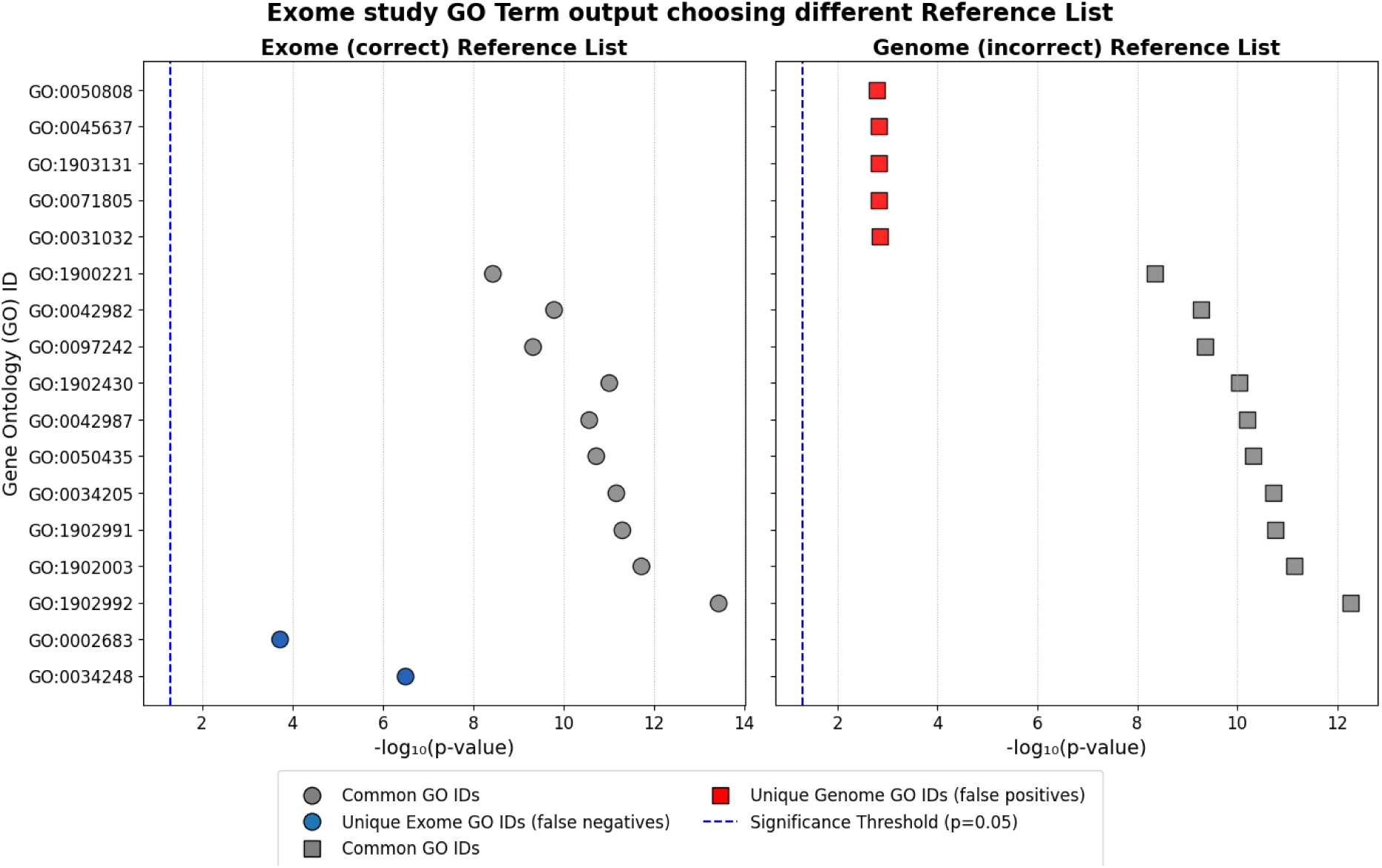
Comparison of GO Biological Process outputs under exome and genome backgrounds for the same datasets. Brain-domain dataset BE1 analyzed using the exome background (left) and the genome background (right). In each panel, each point corresponds to a GO identifier (GO ID) and is positioned by statistical significance (−log10 p-value); circles denote results under the exome background and squares denote results under the genome background. GO ID categories are color-coded as common to both backgrounds (grey circles), unique to the exome background (blue circles; operationally treated here as potential false negatives under a genome background), and unique to the genome background (red squares; operationally treated here as potential false positives under a genome background). The dashed vertical line marks the nominal significance threshold (p = 0.05).

**Table 4.** Most frequently observed GO Biological Process terms by domain and study type. The table summarizes the GO identifiers (GO IDs) most frequently identified across the studies, stratified by biological domain and study type: genome-wide association studies (GWAS; left) and whole-exome sequencing (WES; right). Domains are indicated using the manuscript’s color coding (brain: blue/top; immune: red/middle; metabolism: green/bottom). For each entry, the GO ID and its associated GO Biological Process term are reported.

Top Common GO IDs per Topic: Genome (Left) and Exome (Right)
| **Brain GWAS** |  | **Brain EXOME** |  |
| --- | --- | --- | --- |
| GO ID | GO term | GO ID | GO term |
| GO:0007041 | lysosomal transport | GO:0006869 | Lipid transport |
| GO:0007269 | neurotransmitter secretion | GO:0007611 | learning or memory |
| GO:0045862 | positive regulation of proteolysis | GO:0007613 | memory |
| GO:0051962 | modulation of chemical synaptic transmission | GO:0042391 | regulation of membrane potential |
| GO:0051963 | regulation of calcium ion transport | GO:0050890 | cognition |
| GO:0051968 | regulation of nervous system development |  |  |
| GO:0098815 | regulation of synapse assembly |  |  |
| GO:0099003 | regulation of synaptic transmission |  |  |
| GO:0099504 | modulation of excitatory postsynaptic potential |  |  |
| GO:0099643 | vesicle-mediated transport in synapse |  |  |

| **Immune GWAS** |  | **Immune EXOME** |  |
| --- | --- | --- | --- |
| GO ID | GO term | GO ID | GO term |
| GO:0001819 | regulation of cytokine production | GO:0002696 | positive regulation of leukocyte activation |
| GO:0030098 | lymphocyte differentiation | GO:0007159 | leukocyte cell-cell adhesion |
| GO:0050863 | regulation of T cell activation | GO:0022407 | regulation of cell-cell adhesion |
| GO:1903131 | mononuclear cell differentiation | GO:0022409 | positive regulation of cell-cell adhesion |
| GO:0032944 | regulation of mononuclear cell proliferation | GO:0030098 | lymphocyte differentiation |
| GO:0050670 | regulation of lymphocyte proliferation | GO:0045785 | positive regulation of cell adhesion |
| GO:0070663 | regulation of leukocyte proliferation | GO:0070663 | regulation of leukocyte proliferation |
| GO:0002287 | alpha-beta T cell activation | GO:0007159 | leukocyte cell-cell adhesion |
| GO:0002292 | T cell differentiation | GO:0022407 | regulation of cell-cell adhesion |
| GO:0002293 | alpha-beta T cell differentiation | GO:0051251 | positive regulation of lymphocyte activation |

**Table 4. Most frequently observed GO Biological Process terms by domain and study type.** The table summarizes the GO identifiers (GO IDs) most frequently identified across the studies, stratified by biological domain and study type: genome-wide association studies
| **Metabolism GWAS** |  | **Metabolism EXOME** |  |
| --- | --- | --- | --- |
| GO ID | GO term | GO ID | GO term |
| GO:0006641 | triglyceride metabolic process | GO:0006520 | amino acid metabolic process |
| GO:0034370 | triglyceride-rich lipoprotein particle remodeling | GO:0009063 | amino acid catabolic process |
| GO:0034372 | very-low-density lipoprotein particle remodeling | GO:0016054 | organic acid catabolic process |
| GO:0034375 | high-density lipoprotein particle remodeling | GO:0044282 | small molecule catabolic process |
| GO:0062012 | regulation of small molecule metabolic process | GO:0046395 | carboxylic acid catabolic process |
| GO:0002443 | leukocyte mediated immunity |  |  |
| GO:0002449 | lymphocyte mediated immunity |  |  |
| GO:0002460 | adaptive immune response |  |  |
| GO:0005996 | monosaccharide metabolic process |  |  |
| GO:0006006 | glucose metabolic process |  |  |

Altogether, these results support the simulation-based conclusion that both **biological domain** and **background selection** materially influence the reliability and interpretability of GO Biological Process enrichment outputs.

## Discussion

GO Biological Process overrepresentation analysis is highly sensitive to the choice of background (also referred to as the universe/reference list) (Khatri et al., 2012; Reimand et al., 2019; Wijesooriya et al., 2022; Yu, 2026). Enrichment analysis of GO Biological Processes is widely used to generate a comprehensive view of the results of molecular studies, such as GWAS, WES, and transcriptomics (Ashburner et al., 2000; Carbon et al., 2021; Khatri et al., 2012). These types of analyses, although very useful, are sensitive to bias, such as the choice of background (reference list), annotation, and GO ID sizes and availability of results (Gaudet & Dessimoz, 2017; Haynes et al., 2018; Mubeen et al., 2022). In this study, we focus on quantifying the effect of the choice of inadequate lists in GWAS and WES studies (de Leeuw et al., 2016; Wijesooriya et al., 2022). We show that using an incorrect background can shift enrichment results in both directions, yielding false positives (processes that appear significant only under a mismatched background) and false negatives (processes that are significant under an assay-appropriate background but are missed under an incorrect one) (Cao et al., 2025; Yu, 2026; Ziemann et al., 2024). We further observe that the magnitude and composition of these effects differ between GWAS and WES gene lists and vary across the biological domains considered here (brain, immune, and metabolism) (Haynes et al., 2018; Mubeen et al., 2022).

To identify and summarize background-dependent effects in a reproducible way, we developed a command code (to be made available on GitHub after acceptance) that runs GO enrichment under genome and exome backgrounds for the same input list and partitions significant GO IDs into background-common and background-unique sets (Reimand et al., 2019; Wijesooriya et al., 2022). We focus primarily on GWAS and WES because their intended backgrounds are well defined (genome and exome, respectively), whereas transcriptomic assays do not have a single fixed background (Reimand et al., 2019; Young et al., 2010). For example, microarray analyses should use the set of genes represented by probes on the array, and next-generation sequencing transcriptomic studies depend on library preparation choices (e.g., random primers versus polyA selection) (Reimand et al., 2019; Young et al., 2010). Moreover, gene expression studies typically provide quantitative information that supports more comprehensive approaches than simple overrepresentation testing, however they suffer from non-independence bias between variables. (Ackermann & Strimmer, 2009; Wu & Smyth, 2012; Maleki et al., 2020).

Although the GO website recommends selecting an appropriate background for each analysis, many workflows use the default genome background (Carbon et al., 2021; Reimand et al., 2019; Wijesooriya et al., 2022). This default is generally aligned with GWAS gene lists and with transcriptomic studies derived from whole-genome sampling, but it can penalize analyses based on restricted assays (Yu, 2026; Wijesooriya et al., 2022). For example, a whole-exome sequencing (WES) study only assesses ∼20,000 coding genes; a microarray covers only the genes represented by its probes; an RNA-seq experiment effectively tests only the genes expressed above detection limits (Reimand et al., 2019; Young et al., 2010). In such cases, using all ∼60,000 human genes (including non-coding or unassayed genes) as the background introduces unrelated genes with no chance of appearing in the list (Cao et al., 2025; Yu, 2026). This can produce counterfactual enrichment (Cao et al., 2025; Ziemann et al., 2024). Yu (2026) illustrates this “background bias” with clusterProfiler: using the entire genome vs. a targeted gene set yields different p-values for the same input list (Yu, 2026; Yu et al., 2012; Wu et al., 2021). Intuitively, if the background is bloated with genes that were never detectable in the experiment, then GO categories composed largely of “detectable” (e.g., coding) genes will seem over-represented (Cao et al., 2025; Gaudet & Dessimoz, 2017). As a result, GO terms can falsely appear significant or non-significant purely due to background composition (Cao et al., 2025; Wijesooriya et al., 2022; Ziemann et al., 2024). This is precisely what our study confirms and quantifies.

The effect is most clearly illustrated in Fig. 2, which compares GO enrichment for exome derived input lists when using the genome background versus the exome background. Across GO IDs, the mismatch produces both increased and decreased significance relative to the assay-appropriate background, and the divergence is larger for larger input lists (Fig. 2A versus Fig. 2B) (Mubeen et al., 2022).

The input-size dependence is summarized in Fig. 2C, where both over- and under-significance increase as input size grows (Mubeen et al., 2022). Notably, the trend is not strictly monotone: the proportion of GO IDs that become “more significant” under the genome background shows a break in the range of 50–100 input genes. This behavior is consistent with the fact that the subset of GO IDs that can shift toward greater significance depends on the joint interaction among input size, GO ID size, and how exome genes are distributed across GO IDs, rather than on input size alone (Mubeen et al., 2022). Fig. 6 further illustrates that the distribution of GO ID sizes is not uniform and that differences in the frequency of specific GO ID sizes can affect the aggregate proportion of GO IDs exhibiting inflation (Mubeen et al., 2022).

**Fig. 6.**
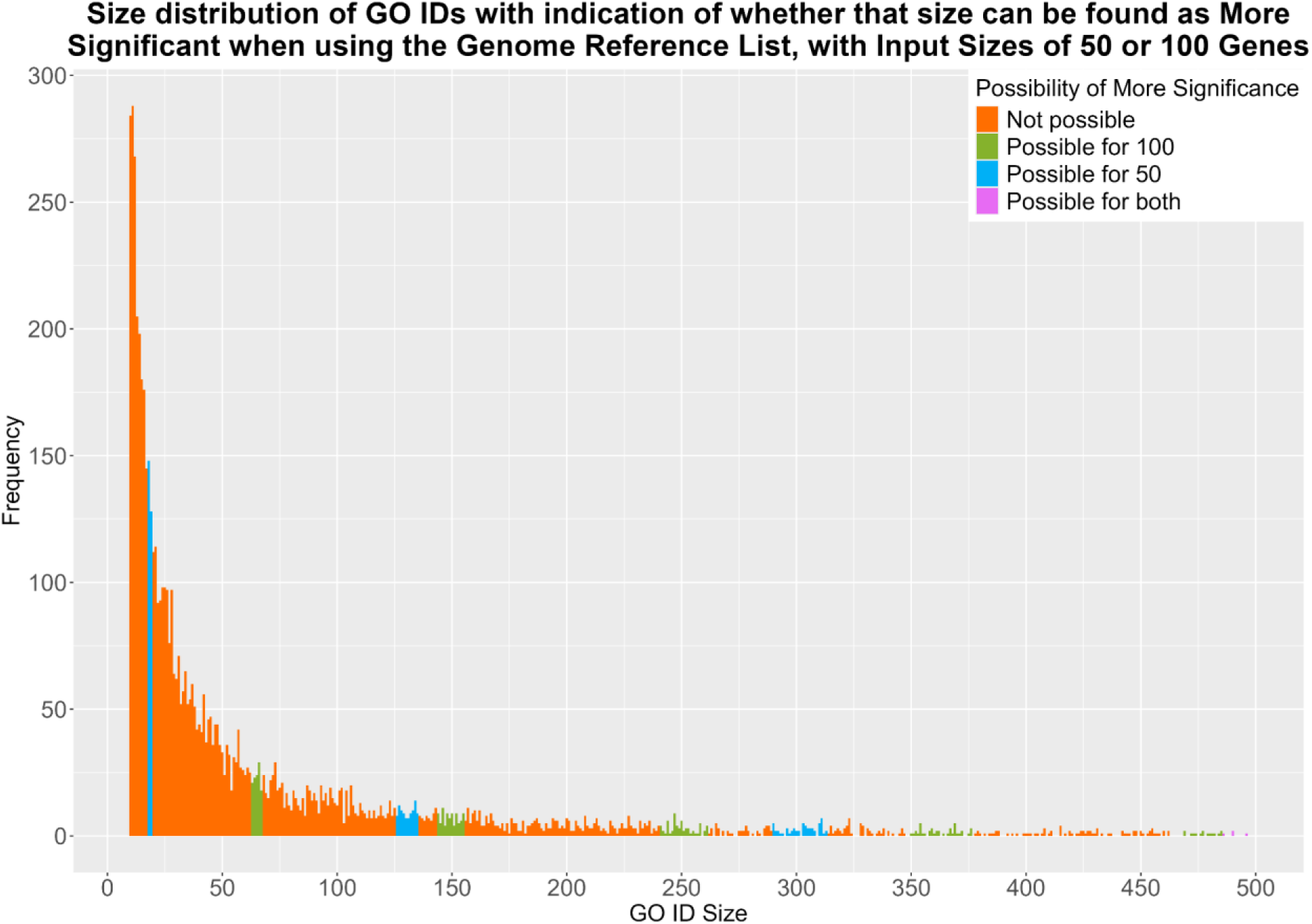
GO ID size distribution and sizes capable of inflation under a genome background for exome-sampled inputs. The figure shows the frequency of GO identifiers (GO IDs) (vertical axis) by GO ID size (horizontal axis) for GO IDs present under both the exome and genome backgrounds. Here, GO ID size is defined as the number of exome-specific genes annotated to the GO ID. Colored bars indicate GO ID sizes that can yield higher significance rates when enrichment is performed using the genome background versus the exome background for input sizes of 50 genes (blue), 100 genes (green), or either 50 or 100 genes (purple). Orange bars indicate GO ID sizes that do not yield higher significance rates under the genome background for input sizes of 50 or 100 genes.

Beyond background mismatch, there are other sources of bias that can affect how genes are associated with GO Biological Processes (Gaudet & Dessimoz, 2017; Khatri et al., 2012). One major problem is limited publication of negative results. (Gaudet & Dessimoz, 2017). A scarcity of reported negative findings can shape the apparent composition of biological processes and reduce the information available to refine functional classification (Gaudet & Dessimoz, 2017).

Another classic issue is “annotation bias”, based on the different number of associations that some genes have with multiple processes, while others have few (Gaudet & Dessimoz, 2017; Haynes et al., 2018). As a result the GO terms that include the “annotation-rich “ genes can be over-enriched even in random lists (Haynes et al., 2018). Conversely, processes involving understudied genes may be under-recognized (Gaudet & Dessimoz, 2017; Haynes et al., 2018). Although there are techniques that try to adjust the annotation bias, there is still an effect of an inadequate reference: when the background includes genes with zero annotations (as in genome vs. exome), those zero-annotation genes dilute the denominator for GO terms comprised of well-annotated genes, leading to inflated significance for those terms (Gaudet & Dessimoz, 2017; Ziemann et al., 2024). In other words, background mismatch is a special case of annotation bias – it effectively changes the proportion of unannotated genes in the “population,” skewing results in favor of densely annotated categories (Gaudet & Dessimoz, 2017; Glass & Girvan, 2014; Ziemann et al., 2024). Ziemann et al. (2024) termed this the “background problem,” noting that some ORA tools automatically drop unannotated genes from the background (Ziemann et al., 2024; Wu et al., 2021). This phenomenon can strongly bias enrichment by shrinking the background to only GO-annotated genes (Ziemann et al., 2024). Our analysis of genome vs. exome backgrounds is directly related to this problem: the genome contains many genes with no GO BP annotations (e.g., non-coding RNAs), whereas the exome background (by construction) contains only protein-coding, mostly GO-annotated genes (Gaudet & Dessimoz, 2017; Ziemann et al., 2024). Using the genome background, we observed exactly what annotation bias predicts: GO categories (comprised mainly of coding genes) appeared more significant than when using the exome background, because the “unannotated” non-coding genes in the genome background made the observed overlap look more unexpected (Gaudet & Dessimoz, 2017; Ziemann et al., 2024). A similar problem results when some genes are under the detection threshold either because they are too short (detectability bias) or because they are low expressed (Young et al., 2010; Mi et al., 2012). In both cases the presence of these genes in the background (i.e. the genome), but not in the actual gene lists, which contain mostly protein-coding genes (the exome) affects the result of the enrichment analysis.

Again, our Monte Carlo simulations confirmed that the wrong background leads to the inflation of GO terms that are predominantly coding (Cao et al., 2025; Mubeen et al., 2022). For instance, we saw GO terms related to neuronal processes or metabolic enzymes (essentially all coding genes) showing up as false enrichments when using a whole-genome reference with exome derived lists, especially as list size grew (Cao et al., 2025; Gaudet & Dessimoz, 2017). By contrast, terms that involve many non-coding elements (few such GO BP terms exist, because it in intrinsic to GO’s nature) would conversely appear less significant than they should (Cao et al., 2025). Figure 2 of our paper illustrates this clearly: points above the diagonal (more significant under genome-background) there are often GO terms in which non-coding genome genes diluted the reference, and the inflation worsened with larger input lists (Cao et al., 2025; Mubeen et al., 2022). The general rule is that we should always match the background to the gene universe of the experiment, and this practice is particularly relevant in experiments of transcriptomics (Reimand et al., 2019; Young et al., 2010).

The observation that genes that are more studied are more “annotation rich” and can therefore bias the enrichment of GO terms, prompted us to investigate the effect of a mismatched universe in different domains, since some areas are more investigated than others (Gaudet & Dessimoz, 2017; Haynes et al., 2018). This unbalanced research focus results in having more enrichment in the more “popular” genes (Haynes et al., 2018). Besides, we need to remember that lack of annotations does not necessarily mean lack of function (Gaudet & Dessimoz, 2017).

To assess whether background effects and potential annotation imbalance differ across domains, we examined gene lists stratified by brain, immune, and metabolism (Haynes et al., 2018). When using the genome background for exome derived inputs (a common misuse), metabolism studies showed fewer background-unique genome GO IDs than the brain and immune domains (Haynes et al., 2018; Wijesooriya et al., 2022). Qualitatively, unique genome GO IDs in metabolism and immune studies tended to remain within their expected domains, whereas the brain domain included some background-unique terms associated with immune processes (Table 2) (Haynes et al., 2018). When examining unique exome GO IDs recovered under the assay-appropriate background (terms that would be missed under the genome background), immune results remained domain-consistent, while brain again included immune and metabolic processes (Table 3) (Haynes et al., 2018). These patterns may reflect true biological interplay and/or differences in annotation density and term structure, but they should be interpreted cautiously (Gaudet & Dessimoz, 2017; Haynes et al., 2018).

We need to use caution in generalizing these results since there are two main limitations:1) Domain assignments were based on broad keyword criteria, and each study was represented by short input lists (20 genes per list), which can increase variability in enrichment outputs (Mubeen et al., 2022); 2), public availability of WES gene lists is limited, reducing the number of exome examples that can be analyzed and increasing uncertainty in domain-level comparisons (Wijesooriya et al., 2022). With the available 6 GWAS and 3 WES studies, the observed domain-related patterns remain suggestive rather than definitive (PMID: 35263338) (Wijesooriya et al., 2022).

Overall, our results show that failing to use an assay-appropriate background in genome and exome studies can both introduce background-specific “significant” processes and mask processes that are significant under the correct background (Cao et al., 2025; Yu, 2026; Wijesooriya et al., 2022). Practically, analyses should (i) select backgrounds that match the assay design, (ii) report the background explicitly, and (iii) perform a sensitivity check by comparing background-common and background-unique GO IDs when plausible alternative backgrounds exist (Reimand et al., 2019; Wijesooriya et al., 2022). More recently, some tools have become available to limit the background issue: clusterProfiler 4.0 has now options to retain the user-specified background (including unannotated genes) (Wu et al., 2021; Ziemann et al., 2024). Nonetheless, the correction of the background is still not common practice, and neither it is to report all the details of molecular and data processing in the studies, which would be essential to adopt the correct conditions in the analysis (Wijesooriya et al., 2022). The provided workflow supports transparent background specification and helps make enrichment-based interpretation more reproducible across studies and pipelines (Reimand et al., 2019; Wijesooriya et al., 2022).

## Acknowledgements

This research was supported by FutureNeuroResearch Ireland Grant No.21/RC/10294_P2, and it was partially supported by SFI (16/IA/4443)

## Declaration of generative AI and AI-assisted technologies in the manuscript preparation process

During the preparation of this work the author(s) used ChatGPT in order to correct the grammar and orthography. After using this tool/service, the author(s) reviewed and edited the content as needed and take(s) full responsibility for the content of the published article.

